# Transcription factor FOXM1 specifies the loading of chromatin DNA to extracellular vesicles

**DOI:** 10.1101/2022.01.27.477315

**Authors:** Yunsheng Zhang, Nana Ding, Yizhen Li, Xiaoyun Zhao, Shiping Yang, Ping Fu, Yousong Peng, Haojie Cheng, Min Ouyang, Ruiping Wang, Yu Wang, Hanyun Liu, Yan Chen, Xiaoqin Huang, Li Yu, Yongjun Tan

## Abstract

Extracellular vesicle DNAs (evDNAs) possess the important diagnostic value for multiple diseases and play roles for horizontally transferring genetic materials among cells. In this study, we have found that transcription factor FOXM1 can mediate the loading of certain chromatin genes or DNA fragments (named FOXM1-chDNAs) to extracellular vesicles (EVs). FOXM1 interacts with LC3 in nucleus and FOXM1-chDNAs (such as *DUX4* gene and Telomere DNA) are specified by FOXM1 and translocated to cytoplasm. These DNAs are released to EVs through the process of an LC3-involved autophagosome-multivesicular body (MVB) transport. The roles of FOXM1 on loading FOXM1-chDNAs to EVs are further confirmed by DNA-FISH experiments, tracing the translocation of selected chromatin loci with the TetO/TetR-GFP method, and PCR analysis of the DNA samples from MVBs and EVs. Furthermore, disrupting the expression of FOXM1 or the process of autophagosome-MVB transport impairs the loading of FOXM1-chDNAs to EVs. This discovery suggests that transcription factor FOXM1 contributes the constitution of evDNAs from nuclear chromatin, providing the first example to explain how chromatin DNA fragments are specified and loaded to EVs. It also provide a foundation to further explore the roles of evDNAs in biological processes such as the horizontal gene transfer.

## Introduction

Extracellular vehicles (EVs) are membranous vesicles released by a variety of cells into the extracellular microenvironment and contain DNAs, RNAs, and proteins acting as intercellular messengers for exchanging variable materials among cells^1,2^. Because extracellular vesicle DNAs (evDNAs) carry the portions of genetic materials from their parent cells under certain physiological or pathological conditions, extensive studies have focused on the diagnostic value of evDNAs for multiple diseases, while their biological functions are not explored as deeply as that of the RNAs or proteins of EVs^3,4^. The evDNAs have been proposed to play a role in so called the horizontal gene transfer (HGT)^5^, which allows to exchange functional genes and DNA fragments among cells^6^. Increasing evidences support that EVs contain functional genes or chromosome DNA fragments as evDNAs and transfer them to recipient cells^7^. For example, LINE-1 retrotransposon as an evDNA of EVs is horizontally transferred among human cancer cells^8^. Another example is that Telomere DNA carried by the EVs from antigen-presenting cells (APCs) is horizontally transferred to primarily naïve T cells and central memory T cells^9^. Therefore, it is critical to understand how chromatin DNA fragments are specified and loaded to EVs, not only for improving the application of evDNAs in diagnosis but also for clarifying their functions in biological processes such as HGT.

Recently, chromatin DNA has been found to translocate to cytoplasm through the interaction between laminB1 and LC3 in nucleus during autophagy^10^, providing a clue to explore the origin of evDNAs. LC3 is a key member of the Atg8 family proteins that are responsible for the cargo selection of autophagy^11^. Besides existing in cytoplasm, LC3 is present in nucleus and interacts with the chromatin-binding laminB1 (a nuclear lamina protein^12^). During autophagy, a part of regular LC3 (LC3-I) need be conjugated with phosphatidyl ethanolamine (PE) to form LC3-PE (LC3-II), which is termed as the lipidation of LC3 and allows the LC3 protein embedding in membranes and then loading cargos to autophagosomes^13^. Although cytoplasmic autophagosomes fuse with endosomes or lysosomes to degrade their components^14,15^, accumulated evidences prove that a portion of endosomes fused with autophagosomes can mature to form multivesicular bodies (MVBs, a special kind of late endosomes^16^) for the secretion of their cargos^1,17,18^. MVBs contain LC3-positive intraluminal vesicles (ILVs)^18^, which can be released to extracellular environment as EVs through the fusion of MVBs with cell membrane^2^. It’s well known that LC3 specifies the loading of proteins and RNAs to EVs^18,19^, confirming that variable materials can be released by EVs through the process of an LC3-involved autophagosome-MVB transport. However, because laminB1 possesses no sequence specificity for its DNA binding^20^, it is hard to figure out how certain chromatin DNA fragments are specified and loaded to EVs, if only basing on the mechanism of laminB1-LC3-mediated chromatin DNA cytoplasmic translocation.

Here, we show a transcription factor-LC3-involved mechanism specifying certain chromatin genes or DNA fragments to EVs during autophagy. FOXM1, a member of Forkhead Box transcription factor family^21^, participates in regulating cell proliferation^22^, DNA damage repair^23^, cell stemness^24^, and metastasis^25,26^ through stimulating gene transcription in nucleus^27^. FOXM1 can specify its binding regions on chromatin directly through its DNA-binding consensus sequence^28^ or indirectly through interacting with other transcription factors such as B-Myb, MuvB, and NFY^29^. In this study, we have confirmed that FOXM1 interacts with LC3 in nucleus and mediates the loading of specific chromatin DNA fragments (named FOXM1-chDNAs) to EVs during autophagy. The interaction between FOXM1 and LC3 is mediated by the LC3-interacting region motif of FOXM1 (317-320aa) and facilitated by the lipidation of LC3, while the FOXM1-LC3 interaction does not disrupt the DNA binding ability of FOXM1. FOXM1-chDNAs, including *DUX4* gene and Telomere DNA, etc., have been identified by analyzing the sequencing data from FOXM1-specific ChIP-seq, LC3-specific ChIP-seq, and evDNA-seq. The loading of specific FOXM1-chDNAs to EVs has been confirmed by DNA-FISH experiments, tracing the translocation of selected chromatin loci with the TetO/TetR-GFP method, and PCR analysis of the DNA samples from MVBs and EVs. We have confirmed that the expression of FOXM1 or the process of LC3-involved autophagosome-MVB transport is essential for the loading of FOXM1-chDNAs to EVs. Together, this study has provided the first example to explain how chromatin DNA fragments are specified and loaded to EVs and a foundation to further explore the roles of evDNAs in biological processes such as HGT.

## Results

### The interaction between FOXM1 and LC3

LC3 mediated the loading of variable materials to EVs through the process of an LC3-involved autophagosome-MVB transport^18,19^. Interestingly, we identified LC3 as a potential partner interacting with transcription factor FOXM1, which directly or indirectly bound to the specific regions on chromatin, from our mass spectrometry analysis of the FOXM1 interactome (data not shown), implicating that FOXM1 might participate in specifying and loading chromatin DNA fragments to EVs. To test this hypothesis, we first confirmed the interaction between FOXM1 and LC3 in nucleus of lung cancer A549 cells. The immunostaining of endogenous FOXM1 and LC3 showed that the two proteins were co-localized in nucleus with the Pearson’s Correlation Coefficient value around 0.66 ± 0.08 (**Fig. 1a**). The co-immunoprecipitation (co-IP) experiments revealed that FOXM1 interacted with LC3, especially with the lipidated LC3-II in nucleus (**Fig. 1b and Extended Data Fig. 1a, b**). A bimolecular fluorescence complementation (BiFC) assay^30^ further confirmed that the FOXM1-LC3 interaction happened mainly in nucleus (**Fig. 1c**). The co-IP of exogenous Flag-FOXM1 with GFP-LC3 WT or G120A mutant, which disrupted the LC3 lipidation on its G120 residue^10^, showed that the deficiency of the LC3 lipidation decreased the FOXM1-LC3 interaction (**Extended Data Fig. 1c, d**), implicating that LC3 embedding in nuclear membranes facilitated its interaction with FOXM1.

**Figure 1.**
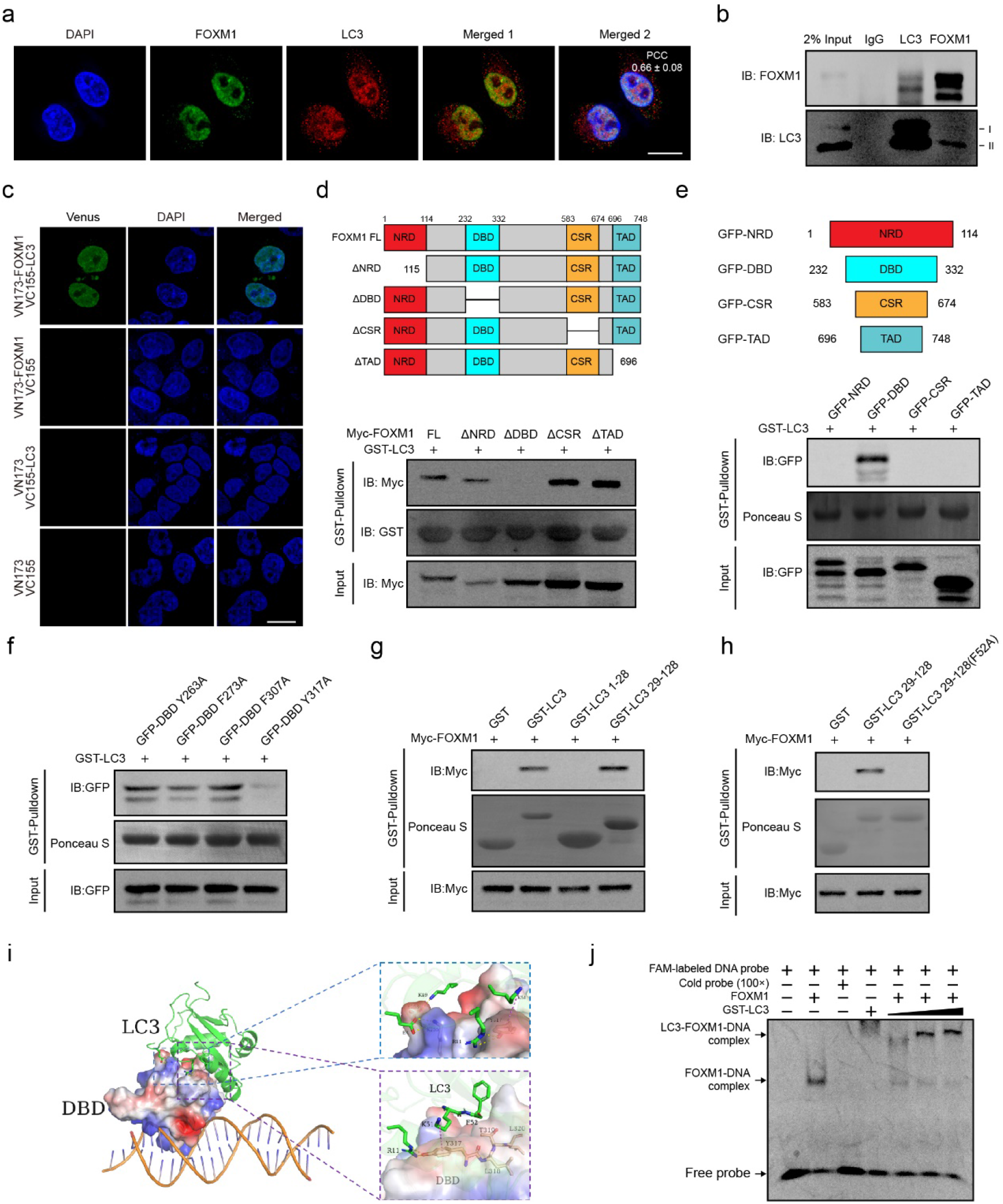
The interaction between FOXM1 and LC3. **(a)** Representative FOXM1 and LC3 images in A549 cells that were fixed by 4% paraformaldehyde and immunostained with anti-FOXM1 and anti-LC3 antibodies. Scale bar, 10 μm. The Pearson Correlation Coefficient (PCC) value of FOXM1 and LC3 colocalization was 0.66±0.08. **(b)** Co-immunoprecipitation of endogenous FOXM1 and LC3 in A549 cells. **(c)** The BiFC assays of FOXM1-LC3 interaction. HEK293T cells were transfected with the indicated combination of split Venus constructs for 24 hrs. Scale bar, 10 μm. **(d)** HEK293T cells (5 x 10^6^) were transfected with Myc-tagged FOXM1 deletion mutant plasmids (5 μg). All cell lysates were pulled down with bacterially purified GST-LC3 protein (30 μg) and subjected to immunoblotting as indicated. **(e)** The GFP-fused FOXM1 truncations were expressed and pulled down with bacterially purified GST-LC3 as described above. **(f)** The GFP-fused FOXM1-DBD point mutations were expressed and pulled down with bacterially purified GST-LC3 as described above. **(g)** The Myc-tagged FOXM1 was pulled down with bacterially purified GST-LC3 truncated mutants as described above. **(h)** The Myc-tagged FOXM1 was expressed and pulled down with bacterially purified GST-LC3 truncated point mutants as described above. **(i)** The molecular docking simulation for FOXM1-LC3 interaction by Rosetta Dock. **(j)** EMSA showed FOXM1, LC3 and DNA formed LC3-FOXM1-DNA complex.

Next, we identified the regions of FOXM1 and LC3 to mediate the FOXM1-LC3 interaction. We screened FOXM1 protein and found that FOXM1 DNA-binding domain (FOXM1-DBD) was required for its LC3 binding (**Fig. 1d**) and FOXM1-DBD interacted with LC3 in GST-LC3-pulldown assays (**Fig. 1e**). Moreover, the overexpression of GFP-FOXM1-DBD abolished the endogenous FOXM1-LC3 interaction (**Extended Data Fig. 1e, f**). It was known that so-called LC3-interacting region (LIR) motif existed in LC3’ substrate proteins^31,32^. We screened the protein sequence of FOXM1-DBD and identified four LIR motifs (263-266, 273-276, 307-310 and 317-320), which were evolutionarily conserved among FOXM1s of human, mouse, rat, and macaque (**Extended Data Fig. 1g**). We generated four FOXM1-DBD mutants (Y263A, F273A, F307A, and Y317A, corresponding to the four LIR motifs respectively), among which only the Y317A mutation on FOXM1-DBD abolished its interaction with LC3 in GST-LC3-pulldown assays (**Fig. 1f**), suggesting that the LIR motif (317-320) in FOXM1-DBD mediated the binding of FOXM1 to LC3. On the other hand, because the residues R10, R11, and F52 of LC3 mediated the binding of LC3 to its protein partners^10,33^, we tested whether the LC3 aa1-28 region (containing R10 and R11) or the LC3 aa29-128 region (containing F52) could interact with FOXM1 in GST-pulldown assays. We found that only the aa29-128 region (containing F52) of LC3 interacted with FOXM1 (**Fig. 1g**), implicating that the residues F52 of LC3 mediated the binding of LC3 to FOXM1. Thus, we generated an LC3 aa29-128 truncated F52A mutant and found that this mutant could not bind to FOXM1 in GST-pulldown assays (**Fig. 1h**). Rosetta Dock^34^ was performed with the known structures of FOXM1-DBD^35^ and LC3^36^ to simulate the FOXM1-LC3 interaction and predicted the formation of hydrogen bonds between the LIR motif (317-320) of FOXM1 and the residue F52 of LC3 (**Fig. 1i**), further supporting the conclusion of the interaction between FOXM1 and LC3 in biochemical experiments.

To test whether the binding of LC3 to FOXM1-DBD affected the DNA-binding ability of FOXM1, we performed the electrophoretic mobility shift assays (EMSAs) with recombinant FOXM1 proteins and a FAM-labeled DNA probe containing putative FOXM1 binding sites. We observed that the DNA probe formed FOXM1-DNA complexes with FOXM1 alone, and created a supershift of LC3-FOXM1-DNA complexes by adding recombinant LC3 proteins in the reactions (**Fig. 1j**), suggesting that the FOXM1-LC3 interaction did not impair the DNA-binding ability of FOXM1 and the LC3-FOXM1-DNA complex could be formed in cells.

### FOXM1-chDNAs are loaded to extracellular vesicles during autophagy

From the results of LC3 or FOXM1 immuno-staining and DNA DAPI-staining of A549 cells, we observed that DNAs, FOXM1, and LC3 co-localized in cytoplasmic vesicles (**Fig. 2a and Extended Data Fig. 2a**), whose numbers were increased by starvation-induced autophagy (from 3% in control cells to 15% in starvation-treated cells) (**Fig. 2b**), suggesting that the formation of LC3-FOXM1-DNA vesicles normally happened in cytoplasm and were enhanced during autophagy. Similar phenomena were also observed from multiple human cancer cell lines, such as Hela and HEK293T cells, or the cells treated with different autophagy-inducing conditions, such as No-amino acid, No-serum, or Rapamycin treatment (20 μM) (data not shown). The BiFC assays with exogenous VN173-FOXM1 and VC155-LC3 further confirmed that FOXM1 and LC3 co-existed in cytoplasmic DNA-containing vesicles (**Fig. 2c**). Interestingly, the expression of RFP-FOXM1 ΔDBD, in which FOXM1-DNA binding domain (DBD) was deleted, could not appear in cytoplasmic DNA-containing vesicles (**Extended Data Fig. 2b**), confirming that the FOXM1 DNA-binding ability was required for the formation of cytoplasmic LC3-FOXM1-DNA vesicles during autophagy. Next, we collected EVs (including Large EVs (lEVs) and Small EVs (sEVs)) from starved A549 cells by serial differential ultracentrifugation^37^ (**Extended Data Fig. 2c**). We showed that both FOXM1 and LC3 existed in the samples of EVs by Western blotting (**Fig. 2d**). Because researchers usually focused on sEVs when studying EVs^38,39^ and only sEVs from our samples possessed EV marker TSG101^40^ (**Fig. 2d**), we just collected sEVs as EV samples in following experiments and characterized their size range around 80-250 nm through nanoparticle tracking analysis and transmission electron microscopy (**Extended Data Fig. 2d and e**). The DNA samples from EVs (evDNAs) were purified and analyzed by the vertical agarose gel electrophoresis (**Fig. 2e**), which showed the size range of evDNAs around 2 to 15 kb (majorly around 12 kb), confirming that DNA, FOXM1, and LC3 co-existed in the samples of EVs.

**Figure 2.**
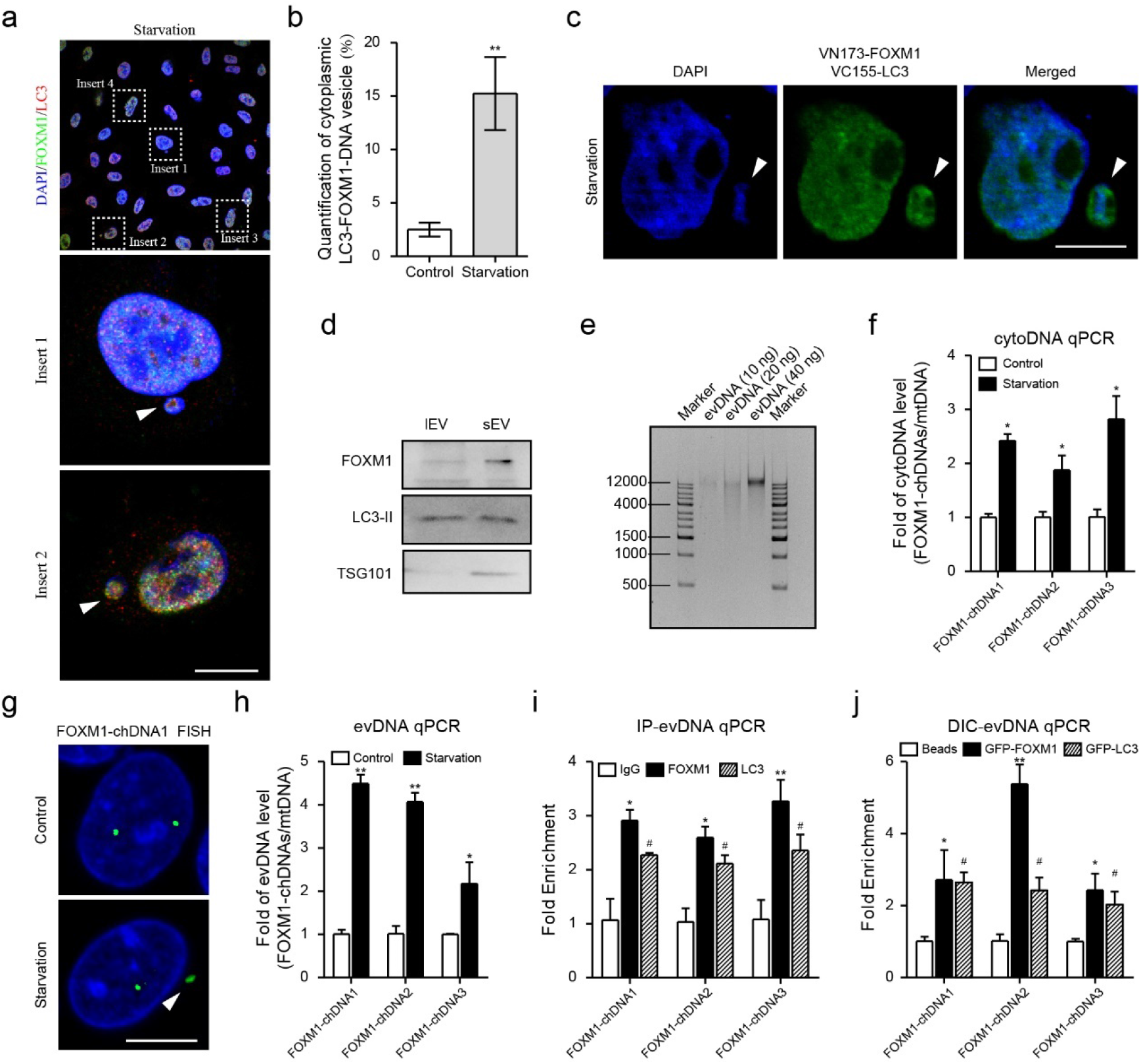
FOXM1-chDNAs are loaded to extracellular vesicles during autophagy. **(a)** Representative LC3-FOXM1-DNA vesicles image in cytoplasm of starved A549 cells (6 hrs) that were immunostained with anti-FOXM1 and anti-LC3 antibodies. The white arrows indicated LC3-FOXM1-DNA vesicle. Scale bar, 10 μm. **(b)** Quantification of cytoplasmic LC3-FOXM1-DNA vesicle in non-starved or starved A549 cells (6 hrs). (n = 4 independent experiments). \*\**P*<0.01; unpaired two-tailed. **(c)** The BiFC assays showed that the exogenous VN173-FOXM1, VC155-LC3 and chromatin DNA vesiculated in cytoplasm of starved A549 cells (6 hrs). Scale bar, 10 μm. **(d)** EVs were collected from starved A549 cells (48 hrs) and subjected to immunoblotting for TSG101, FOXM1 and LC3. **(e)** evDNAs were purified from EVs collected from starved A549 cells (48 hrs). The different concentrations of evDNAs were analyzed by 1% vertical agarose gel electrophoresis. **(f)** The cytoplasmic DNA (cytoDNA) was extracted from non-starved or starved A549 cells (24 hrs) for qPCR. The mitochondria DNA (mtDNA, *ND*-2 gene) was used as the internal loading control. The Bars, mean±s.e.m.; n=3; \**P*<0.05; NS, non-significant; unpaired two-tailed. **(g)** Representative DNA-FISH images of FOXM1-chDNA1-adjacent region (10 Kb probe) in starved A549 cells (6 hrs). The white arrows indicated FOXM1-chDNA1. **(h)** The evDNA was extracted from non-starved or starved A549 cells (48 hrs) for qPCR. The data were normalized by mtDNA. **(i)** The qPCR analyses of FOXM1-chDNAs in anti-FOXM1-IP evDNA or anti-LC3-IP evDNA from EVs collected from starved A549 cells (48 hrs). **(j)** The qPCR analyses of FOXM1-chDNAs in the DIC-purified GFP-FOXM1-specific EVs or DIC-purified GFP-LC3-specific EVs collected from starved A549 stably expressing GFP-FOXM1 or GFP-LC3 cells (48 hrs). The Bars, mean±s.e.m.; n=3; \**P*<0.05, #*P*<0.05, \*\**P*<0.01; unpaired two-tailed.

To identify the chromatin DNA fragments specified by FOXM1 (FOXM1-chDNAs) that were finally loaded to EVs, we first performed chromatin immunoprecipitation sequencing (ChIP-seq) analysis for FOXM1 or LC3 separately with starved A549 cells and obtained 103 specific loci on chromatin bound by both FOXM1 and LC3 (**Table S**) from the overlapping of FOXM1-ChIP fragments (n=8, 838) (GSE216672) and LC3-ChIP fragments (n=7, 951) (GSE216672). Then we performed evDNA-sequencing analysis with the EVs of starved A549 cells and obtained 15, 544 evDNAs (GSE216672). After analyzing the overlapped sequences of FOXM1-LC3 ChIP loci and evDNAs, we identified 25 chromatin loci as FOXM1-chDNAs (**Table**), which could be annotated as functional genes (n=8, including *DUX4* gene), telomere (n=1), centromere (n=8), and nonsense regions (n=8). We chose three FOXM1-chDNAs (FOXM1-chDNA1 from a telomere locus, FOXM1-chDNA2 from a nonsense region, and FOXM1-chDNA3 from *DUX4* locus) as examples to perform ChIP-qPCR and confirmed the binding of FOXM1 and LC3 on these FOXM1-chDNAs at chromatin but not on the control region such as β-actin promoter (**Extended Data Fig. 2f, g**).

**Table.** FOXM1-chDNAs.

| Names | Sequence signatures | Chr | geneID |
| --- | --- | --- | --- |
| FOXM1-chDNA1 | Telomere | 18 | ENSG00000263006 |
| FOXM1-chDNA2 | Nonsense region | 9 | ENSG00000228522 |
| FOXM1-chDNA3 | Coding region (*DUX4*) | 4 | ENSG00000260596 |
| FOXM1-chDNA4 | Coding region (*PGK1*) | X | ENSG00000102144 |
| FOXM1-chDNA5 | Coding region (*ITFG2*) | 12 | ENSG00000111203 |
| FOXM1-chDNA6 | Coding region (*FOXM1*) | 12 | ENSG00000111206 |
| FOXM1-chDNA7 | Coding region (*FOXM1*) | 12 | ENSG00000111206 |
| FOXM1-chDNA8 | Coding region (*TBC1D5*) | 3 | ENSG00000131374 |
| FOXM1-chDNA9 | Coding region (*XR_946063.2*) | 10 | ENSG00000166317 |
| FOXM1-chDNA10 | Coding region (*XR_943570.1*) | 14 | ENSG00000258314 |
| FOXM1-chDNA11 | Centromere | 5 | ENSG00000170571 |
| FOXM1-chDNA12 | Centromere | 1 | ENSG00000224857 |
| FOXM1-chDNA13 | Centromere | 20 | ENSG00000227195 |
| FOXM1-chDNA14 | Centromere | 3 | ENSG00000251727 |
| FOXM1-chDNA15 | Nonsense region | 16 | ENSG00000261239 |
| FOXM1-chDNA16 | Nonsense region | 16 | ENSG00000261239 |
| FOXM1-chDNA17 | Nonsense region | 16 | ENSG00000261239 |
| FOXM1-chDNA18 | Nonsense region | 16 | ENSG00000261239 |
| FOXM1-chDNA19 | Nonsense region | 16 | ENSG00000261239 |
| FOXM1-chDNA20 | Centromere | 17 | ENSG00000263433 |
| FOXM1-chDNA21 | Centromere | 17 | ENSG00000263433 |
| FOXM1-chDNA22 | Centromere | 17 | ENSG00000263433 |
| FOXM1-chDNA23 | Centromere | 17 | ENSG00000264970 |
| FOXM1-chDNA24 | Nonsense region | 1 | ENSG00000276118 |
| FOXM1-chDNA25 | Nonsense region | 5 | ENSG00000284042 |

To verify that FOXM1-chDNAs were translocated to cytoplasm and finally loaded to EVs, we first purified cytoplasmic DNA (cytoDNA) particularly^41^ and found the levels of the three FOXM1-chDNAs were elevated in cytoplasm of starved A549 cells (**Fig. 2f**). A DNA-FISH probe (10 Kb) specific for the FOXM1-chDNA1-adjacent region was generated and DNA-FISH experiments showed that FOXM1-chDNA1 appeared in cytoplasm post starvation (**Fig. 2g**), confirming that FOXM1-chDNAs were translocated to cytoplasm during autophagy. Then, we purified evDNAs from the EVs of starved A549 cells and found the levels of the three FOXM1-chDNAs were elevated in the EVs (**Fig. 2h**), confirming that FOXM1-chDNAs were finally loaded to EVs during autophagy. In addition, we also treated the EVs of starved A549 cells with the Plasmid-Safe ATP-dependent DNase (PS), which only digested linear DNA^42^, and observed the decreased levels of the three FOXM1-chDNAs from the PS-treated samples (**Extended Data Fig. 2h**), identifying that FOXM1-chDNAs existed as linear DNA on outside the lumen of EVs. This result was consistent with the previous finding that evDNAs mainly localized on the outside the lumen of EVs^43^. Then, we treated the EVs of starved A549 cells with formaldehyde to cross-link their DNA and proteins and the lysates of EVs were immune-precipitated with anti-FOXM1 or anti-LC3 antibodies for evDNA purification. The levels of the three FOXM1-chDNAs in the FOXM1 or LC3 specific IP-evDNA samples were significantly elevated (**Fig. 2i**), further confirming that FOXM1-chDNAs were bound by FOXM1 or LC3 on EVs. We also collected FOXM1-specific EVs or LC3-specific EVs from GFP-FOXM1 or GFP-LC3-overexpressed A549 cells through the direct immunoaffinity capture (DIC) approach^1^ with GFP-Trap and observed the elevated levels of the three FOXM1-chDNAs in the DIC samples (**Fig. 2j**), suggesting that both exogenous GFP-FOXM1 and GFP-LC3 facilitated the loading of FOXM1-chDNAs to EVs. Together, these data suggested that FOXM1 and LC3 mediated the loading of FOXM1-chDNAs to EVs during autophagy.

### The FOXM1-chDNAs knocked-in with the TetO array were loaded to EVs

To confirm the translocation of FOXM1-chDNAs to EVs, we knocked in a TetO array (96x) at the loci of selected FOXM1-chDNAs by CRISPR/cas9 in A549 cells (**Extended Fig. 3a**). DNAs containing the TetO array then were visualized directly by exogenously expressed TetR-EGFP in cells^44^. Because Telomere DNA was proved to load to EVs^9^, we first edited the adjacent region of FOXM1-chDNA1 (a Telomere locus) at chromosome 18 (**Extended Fig. 3b**) and observed the signal of TetO-Telomere in nucleus followed by its elevated levels in cytoplasm post starvation (**Extended Fig. 3c, d**), confirming the validity of the TetO/TetR-GFP method. Next, we edited the locus of FOXM1-chDNA3 (*DUX4* locus) at chromosome 4 (**Extended Fig. 3b**) for following experiments. DUX4 was identified as a transcription factor playing roles in embryonic development^45^ and in promoting immune evasion of cancers^46^. The signals of TetO array-containing DUX4 DNA (TetO-DUX4) could be visualized in nucleus of A549*^TetO-DUX4^* cells and its cytoplasmic signals were induced by starvation (**Fig. 3a**), which were further confirmed by the qPCR results of the cytoDNA of TetO-DUX4 (**Fig. 3b**). The signal of TetO-DUX4 observed in cytoplasm post starvation was co-localized with the immunofluorescent signal of LC3 (**Fig. 3c**), confirming that FOXM1-chDNA3 (DUX4 locus) appeared in autophagosomes during autophagy.

**Figure 3.**
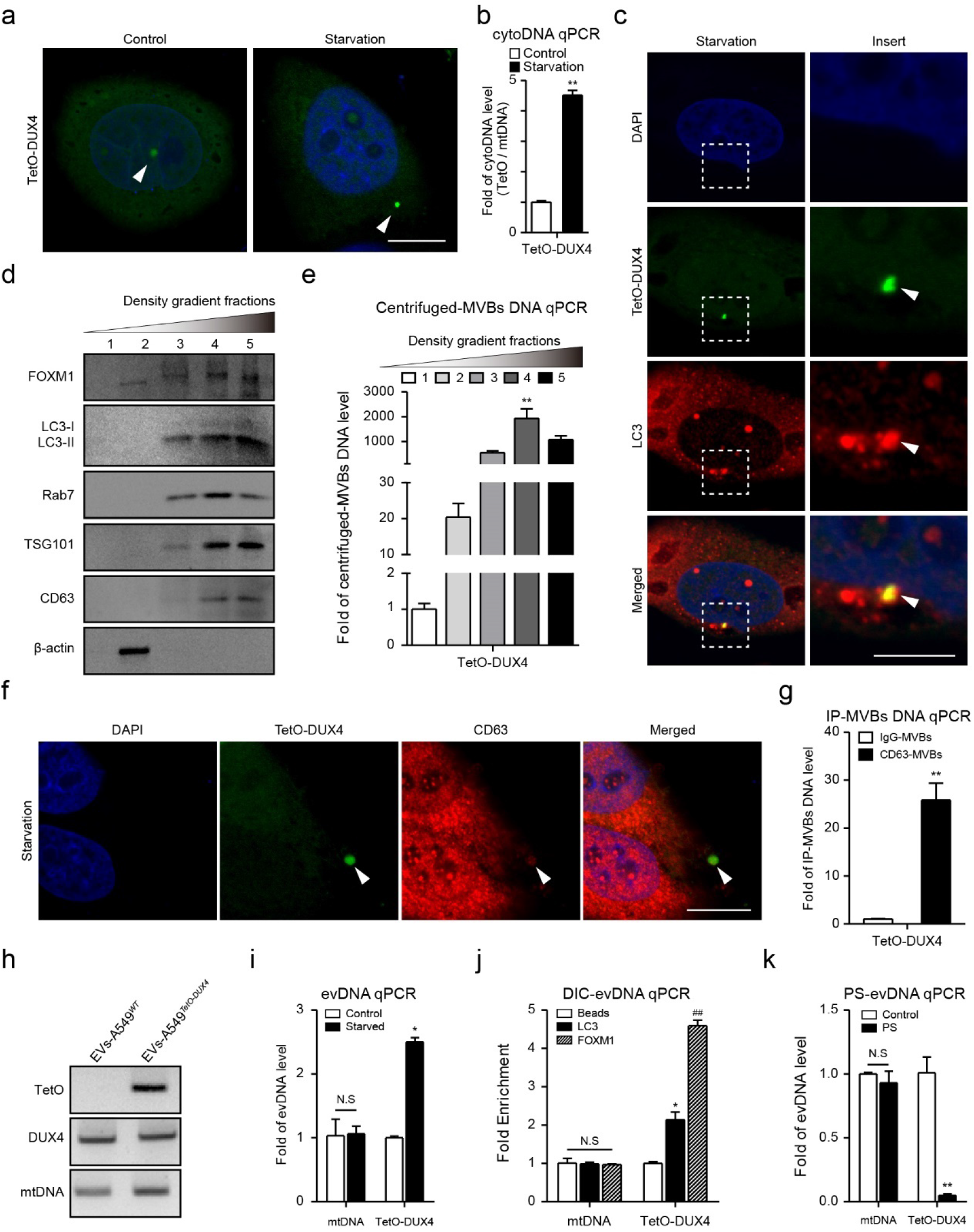
The FOXM1-chDNAs knocked-in with the TetO array were loaded to EVs. **(a)** Representative TetO-DUX4 images in starved A549*^TetO-DUX4^* cells (6 hrs) that were transiently expressed TetR-EGFP. The white arrows indicated TetO-DUX4. Scale bar, 10 μm. **(b)** The A549*^TetO-DUX4^* cells were treated with starvation and cytoplasmic DNA (cytoDNA) was extracted 24 hours later for qPCR. The mtDNA was internal control. The Bars, mean±s.e.m.; n=3; \*\**P*<0.01; NS, non-significant; unpaired two-tailed. **(c)** Immunostaining of the endogenous LC3 and TetO-DUX4 in starved A549*^TetO-DUX4^* cells (6 hrs) that were exogenously expressed TetR-EGFP. The white arrows indicated TetO-DUX4 that was translocated into cytoplasmic autophagosome. Scale bar, 10 μm. **(d)** MVB compartments were isolated from starved A549*^TetO-DUX4^* cells (24 hrs) by discontinuous sucrose gradient and subjected to immunoblotting. **(e)** The qPCR analyses of TetO-DUX4 in MVB fractions collected from starved A549*^TetO-DUX4^* cells (24 hrs) by discontinuous sucrose gradient. The Bars, mean±s.e.m.; n=3; \*\**P*<0.01; NS, non-significant; unpaired two-tailed. **(f)** Immunostaining of the endogenous CD63 and TetO-DUX4 in starved-A549*^TetO-DUX4^* cells (6 hrs). The white arrows indicated TetO-DUX4 that was translocated into cytoplasmic MVB. Scale bar, 10 μm. **(g)** The qPCR analyses of TetO-DUX4 in MVBs collected from starved-A549*^TetO-DUX4^* cells (24 hrs) by CD63-immunoprecipitation. The Bars, mean±s.e.m.; n=3; \*\**P*<0.01; NS, non-significant; unpaired two-tailed. **(h)** PCR was used to amplify the TetO-DUX4, DUX4, and mitochondria DNAs in the evDNAs collected from starved A549*^WT^* or A549*^TetO-DUX4^* cells (48 hrs). **(i)** The qPCR analyses of TetO-DUX4 in the EVs collected from starved-A549*^TetO-DUX4^* cells (48 hrs). **(j)** The qPCR analyses of TetO-DUX4 in the DIC-purified FOXM1-specific EVs or LC3-specific EVs collected from starved-A549*^TetO-DUX4^* cells (48 hrs). **(k)** The qPCR analyses of TetO-DUX4 in the Plasmid-Safe ATP-dependent DNase (PS)-digested EVs collected from starved-A549*^TetO-DUX4^* cells. The Bars, mean±s.e.m.; n=3; \**P*<0.05; #*P*<0.05; \*\**P*<0.01; ##*P*<0.01; unpaired two-tailed.

Because autophagosomes could fuse with endosomes to mature to MVBs^16^ that finally released their cargos as EVs through fusing with cell membrane^2^, we intended to test whether cytoplasmic TetO-DUX4 appeared in MVBs. We used the method of discontinuous sucrose gradient to prepare different fractions to define MVB compartments^47^ from postnuclear supernatants of A549*^TetO-DUX4^* cells. Based on the MVB specific markers, such as Rab7^48^, CD63^49^, or TSG101^50^, we first confirmed that FOXM1, same as LC3, existed in the MVB fractions (**Fig. 3d**). The DNA samples were prepared from the harvested fractions and the results of qPCR specific for TetO-DUX4 DNA showed its significant enrichment in the MVB fractions (**Fig. 3e**). Furthermore, we observed that the signal of TetO-DUX4 in cytoplasm post starvation was co-localized with the immunofluorescent signal of CD63 (**Fig. 3f**). We also isolated the samples of MVBs by immunoprecipitating A549*^TetO-DUX4^* cell lysates with anti-CD63 antibodies^51^ to prepare their DNA, which showed the elevated levels of TetO-DUX4 DNA in these MVBs compared to the IgG-IP controls (**Fig. 3g**), suggesting that this FOXM1-chDNA was transported to MVBs during autophagy.

We collected the EVs from A549 cells and A549*^TetO-DUX4^* cells to prepare DNA samples and the results of PCR confirmed that TetO-DUX4 DNA appeared in EVs (**Fig. 3h**). The starvation in A549*^TetO-DUX4^* cells resulted in the elevated levels of TetO-DUX4 DNA in EVs (**Fig. 3i**). We also collected EVs from A549*^TetO-DUX4^* cells through the DIC approach with anti-FOXM1 or anti-LC3 antibodies to prepare DNA samples and observed the elevated levels of TetO-DUX4 DNA in the samples of FOXM1-specific EVs or LC3-specific EVs (**Fig. 3j**), suggesting that FOXM1 and LC3 were involved in loading TetO-DUX4 DNA to EVs. Moreover, after the EVs were treated with PS DNase, the decreased levels of TetO-DUX4 DNA were observed but not for the mitochondrial DNA controls (**Fig. 3k**), implicating that TetO-DUX4 DNA existed as linear DNA on the outer membrane of EVs. Together, we confirmed that this FOXM1-chDNA was loaded to EVs during autophagy.

### The loading of FOXM1-chDNAs to EVs relied on FOXM1 and the autophagosome-MVB process

We found that the PCR-amplified DNA samples from *DUX4* locus were able to pulled down FOXM1 or lipidated LC3-II *in vitro* (**Fig. 4a**), proving the two proteins binding to this FOXM1-chDNA. To confirm the roles of FOXM1 on the loading of FOXM1-chDNAs to EVs, we generated an A549^*FOXM1*-/-^ cell line by the CRISPR/cas9 technology targeting *FOXM1* gene in A549 cells (**Extended Data Fig. 4a**). In A549^*FOXM1*-/-^ cells, the levels of all the tested three FOXM1-chDNAs (FOXM1-chDNA1 from a telomere locus, FOXM1-chDNA2 from a nonsense region, and FOXM1-chDNA3 from *DUX4* locus) were decreased in cytoplasm compared to that of wild type control cells post starvation in the qPCR analysis of purified cytoDNA samples (**Fig. 4b**). Consequently, the EVs from A549^*FOJXM1*-/-^ cells contained much lower levels of the three FOXM1-chDNAs than that from control cells post starvation (**Fig. 4c**). Furthermore, we knocked down the expression of FOXM1 in A549*^TetO-DUX4^* cells by FOXM1 siRNA (**Extended Data Fig. 4b**) and observed that the cytoplasmic levels of TetO-DUX4 signals were decreased in the cells post starvation (**Fig. 4d**). In addition, we constructed a linear DNA containing 72x FOXM1 binding sites (M1-binding DNA) that was bound by recombinant FOXM1 *in vitro* (**Extended Data Fig. 4c**) or FOXM1-DBD-GFP protein *in vivo* when transfected into cells (**Extended Data Fig. 4d**). We transfected this M1-binding DNA or control DNA at the equal amount into cells and the qPCR analysis of the isolated evDNA samples showed that the levels of M1-binding DNA in EVs were higher than that of control DNA (**Fig. 4e**), suggesting that FOXM1 facilitated the FOXM1-recognized DNA fragments transporting to EVs. We also transfected M1-binding DNA into A549^*FOXM1*-/-^ cells and found that the levels of M1-binding DNA in the EVs of A549^*FOXM1*-/-^ cells were significantly lower than that of wild type control cells (**Fig. 4f**). Together, the data suggested that the loading of FOXM1-chDNAs to EVs relied on FOXM1 in cells.

**Figure 4.**
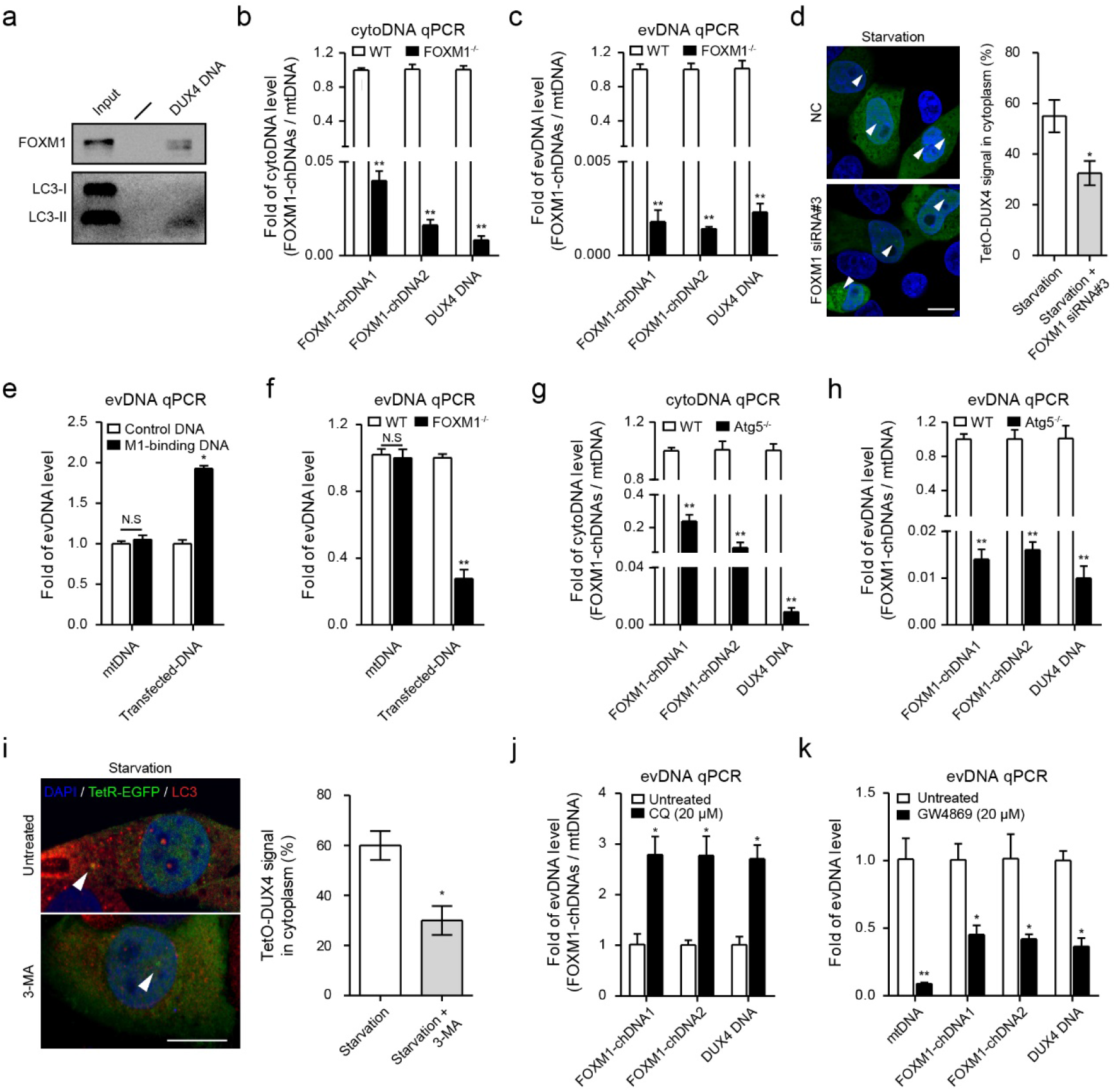
The loading of FOXM1-chDNAs to EVs relied on FOXM1 and the autophagosome-MVB process. **(a)** A549 cell lysates were incubated with biotinylated DUX4 DNA (500 ng). The bound proteins were immunoprecipitated with streptavidin microbeads and blotted by anti-FOXM1 or LC3 antibodies. **(b, c)** The qPCR analyses of three FOXM1-chDNAs in the cytoplasm or EVs collected from starved A549 or A549^*FOXM1*-/-^ cells. The data were normalized by mtDNA. The Bars, mean±s.e.m.; n=3; \*\**P*<0.01; unpaired two-tailed. **(d)** A549*^TetO-DUX4^* cells were transfected with 50 nM *FOXM1* siRNA and 1 μg TetR-EGFP plasmid. After the transfection (48 hrs), cells were starved (6 hrs) and imaged for the signal of TetO-DUX4 (left). The white arrows indicated TetO-DUX4. Scale bar, 10 μm. The TetO-DUX4 signal in cytoplasm was quantitated (right). n = 3 independent experiments; \**P*<0.05; unpaired two-tailed. **(e)** HEK293T cells were transfected with linear M1-binding or control DNA at the equal amount. After the transfection (24 hrs), all cells were starved (48 hrs) for collecting EVs. The qPCR analyses were performed for the mtDNA and transfected DNA in evDNAs purified from the EVs. **(f)** A549 and A549^*FOXM1*-/-^ cells were transfected with linear M1-binding DNA at the equal amount. After the transfection (24 hrs), all cells were starved (48 hrs) for collecting EVs. The qPCR analyses were performed for the mtDNA and transfected DNA in evDNAs purified from the EVs. The Bars, mean±s.e.m.; n=3; \**P* < 0.05; \*\**P*<0.01; NS, non-significant; unpaired two-tailed. **(g, h)** The qPCR analyses of three FOXM1-chDNAs in the cytoplasm or EVs collected from starved A549 or A549^*Atg5*-/-^ cells. The data were normalized by mtDNA. The Bars, mean±s.e.m.; n=3; \*\**P*<0.01; unpaired two-tailed. **(i)** A549*^TetO-DUX4^* cells were transfected with TetR-EGFP plasmid (1 μg). After the transfection (24 hrs), cells were treated with 3-MA (5 mM) and starved (6 hrs). Cells were immunostained with anti-LC3 antibodies and imaged for TetO-DUX4. The white arrows indicated the signal of TetO-DUX4. Scale bar, 10 μm. The TetO-DUX4 signal in cytoplasm was compared (right). n = 3 independent experiments; \**P*<0.05; unpaired two-tailed. **(j)** EVs were collected from A549 cells that were treated with CQ (20 μM) and starvation (24 hrs). The qPCR analyses were performed for the three FOXM1-chDNAs in evDNAs purified form the EVs. **(k)** EVs were collected from A549 cells that were treated with GW4869 (20 μM) and starvation (24 hrs). The qPCR analyses were performed for the three FOXM1-chDNAs in evDNAs purified form the EVs.

To confirm the formation of autophagosomes contributing the loading of FOXM1-chDNAs to EVs, we generated an A549^*Atg5*-/-^ cell line by the CRISPR/cas9 technology targeting *Atg5* gene in A549 cells (**Extended Data Fig. 4e**). It was well known that the Atg5 knockout resulted in the losing of LC3 lipidation and abolished the formation of autophagosomes^52,53^. In A549^*Atg5*-/-^ cells, the cytoplasmic and EVs’ levels of the three FOXM1-chDNAs were lower than that of control cells post starvation (**Fig. 4g, h**), correlated with the decreased levels of FOXM1 and LC3 in the EVs from A549^*Atg5*-/-^ cells (**Extended Data Fig. 4f**). In addition, we treated A549*^TetO-DUX4^* cells with 3-Methyladenine (3-MA), which prevented the formation of autophagosomes^54^, and observed that the cytoplasmic levels of TetO-DUX4 signals were also decreased in the cells post starvation (**Fig. 4i**). To confirm the formation of MVBs contributing the loading of FOXM1-chDNAs to EVs, we treated A549 cells with chloroquine (CQ), a lysosome inhibitor that facilitated the formation of MVBs by inhibiting the fusion of MVBs and lysosomes^16,55^. We observed that the levels of the three FOXM1-chDNAs were elevated in the EVs of the CQ-treated cells post starvation (**Fig. 4j**). In addition, we treated A549 cells with a chemical compound GW4869, a neutral sphingomyelinase inhibitor that prevented the fusion of MVBs with cell membrane and decreased the secretion of EVs^56^. We collected the EVs from the equal numbers of A549 cells non-treated or treated with GW4869 and found that the secretion of TSG101 via EVs was decreased by the GW4869 treatment, confirming the abolished MVB-cell membrane fusion (**Extended Data Fig. 4g**). Consequently, the secretion of FOXM1 or LC3 via EVs was also decreased by the GW4869 treatment (**Extended Data Fig. 4g**). Through the qPCR analysis of the prepared evDNA samples, we observed that the levels of the three FOXM1-chDNAs in EVs, similar as that of mtDNA, were decreased by the GW4869 treatment, suggesting that the secretion of FOXM1-chDNAs to EVs were majorly mediated by MVBs (**Fig. 4k**). Together, the data suggested that the loading of FOXM1-chDNAs to EVs relied on the autophagosome-MVB process.

## Discussion

In this study, we have discovered that transcription factor FOXM1 mediates the loading of chromatin DNA fragments to EVs. Specifically, certain FOXM1-bound chromatin DNA fragments (FOXM1-chDNAs) form LC3-FOXM1-DNA vesicles with FOXM1 and LC3 in cytoplasm during autophagy. The cytoplasmic LC3-FOXM1-DNAs are released to EVs through the process of an autophagosome-MVB transport, providing an example of the transcription factor determining the sources of evDNA from chromatin DNA (**Extended Data Fig. 5**). FOXM1 can theoretically specify its binding regions in chromatin, providing a clue to determine the sources of evDNA from certain chromatin DNA loci. Based on this understanding, the accuracy and specificity of disease diagnosis can be improved in the future by focusing on specific sequences from patient evDNAs.

Although this study focuses on FOXM1, we anticipate that the FOXM1-LC3 interaction is not an isolated event in nucleus. We have found that other Forkhead Box transcription factors such as FOXA2 and FOXP2 can interact with LC3 (data not shown), implicating that these two transcription factors may also participate in determining the sources of evDNA from chromatin DNA. In addition, LC3 interacts with multiple transcription factors that exist in EVs, based on the results of mass spectrometry analysis^18,37^, suggesting that these transcription factors may also play a similar role as FOXM1 for specifying evDNA from chromatin DNA. Transcription factors are generally considered to bind to specific DNA regions of chromosome and regulate specific gene expression in nucleus^57^. The findings of this study expand the understanding of the functions of transcription factors and provide a new perspective to explore their roles in regulating the composition of EVs’ DNA components. Furthermore, the further enrichment analysis of the specific sequences of patient evDNAs can predict certain transcription factors that bind to the sequences, so as to establish the association between the transcription factors and diseases to deduce the pathogenesis or disease progression.

The FOXM1-chDNAs identified in this study can be annotated as functional genes, telomere, centromere, and nonsense regions, implicating their potential effects in the cells that receive these DNAs. Interestingly, a recent study confirmed that Telomere DNAs carried by the EVs from antigen-presenting cells can horizontally transfer to T cells and promote long-term immunological memory by rescuing T cells from senescence^9^. We have observed the signals of TetO-DUX4 in recipient cells that are treated with the A549*^TetO-DUX4^*-derieved EVs and noticed the changes of phenotypes of recipient cells that are transfected with the PCR-amplified DNA of *DUX4* locus (data not shown), providing another example that the certain FOXM1-chDNAs of donor cells can perform specific biological functions in recipient cells after the horizontal gene transfer (HGT) and may regulate the extracellular microenvironment of the cells. Furthermore, because the EVs of local cells are able to enter the circulatory system, we speculate that certain FOXM1-chDNAs may possess relevant roles in HGT to distant tissues. These hypotheses are being investigated in our laboratory.

FOXM1 is generally considered as a transcription factor regulating gene expression in nucleus of cells^27^. This study extends its function to loading certain chromatin DNA fragments to EVs. One key question still remaining is how chromatin DNA breaks to specific fragments. So far, the appearance of chromatin DNA in cytoplasm is mediated mainly by the micronuclei and cytoplasmic chromatin fragment pathways^58^, while both the mechanisms cannot explain the specific breakage of chromatin DNA. Although nuclear LC3 interacts with LaminB1 to result in the cytoplasmic translocation of transcriptionally inactive heterochromatin DNA^10^, the mechanism of specific DNA fragments breaking off chromosomes has not been explored. Recently, the complex of Rec114, Mei4 and Mer2 in cells has been reported to control the precise DNA breakage^59^, suggesting that the breakage of chromatin can be precisely achieved. Future studies of FOXM1’s interaction with the controlled DNA breakage machinery will reveal how relevant chromatin DNA fragments are precisely produced.

## Methods

### The complete Materials and Methods were described in Supplementary Materials

#### ChIP sequencing, evDNA sequencing, Bioinformatics analysis, RT-qPCR

For sequencing, ChIP and evDNA samples were prepared as described previously^60^. The ChIP and evDNA were purified and used for constructing sequencing libraries with a NEBNext Ultra DNA Library Prep Kit for Illumina (New England Biolabs, Ipswich, MA, USA). The library quantifications were assessed on the Agilent Bioanalyzer 2100 system. The library preparations were sequenced on an Illumina Hiseq platform and 50 bp single-end reads or 150 bp paired-end reads were generated.

For bioinformatics analysis, the following pipeline was used for analysis of all sequence data sets. Firstly, fastp (version 0.20.0, parameter: ‘-q 15 -u 40 -n 6 -l 15’)^61^ was used to trim adaptors and low-quality reads. Then, we evaluated the quality of NGS short reads by searching them on the FastQC (version 0.11.8) program (http://www.bioinformatics.babraham.ac.uk/projects/fastqc). Thirdly, the remaining reads were aligned to human genome hg38 by bwa men (version 0.7.17,parameter: ‘-t 20’)^62^, since ChIP-Seq and evDNA-Seq interrupted fragments were usually small and the percentage of unique sequences in the total number of sequences was the focus of attention. Finally, peak calling of the aligned reads was made using macs3 (version 3.0.0a6, parameter: ‘-f BAM -g hs -p 0.1 -B’)^63^, and peaks were annotated using the R/Bioconductor package ChIPseeker^64^. Browser views of peaks are shown using Integrated Genomics Viewer (IGV; broadinstitute.org/igv)^65^.

For qPCR, the following primers were used for qPCR analyses of chromatin loci. FOXM1-chDNA1 (chr18): forward, 5’-TGA TCA CCC AGG AGA CGG-3’; reverse, 5’-CTT GGG TGA TCA GTG GCG AG-3’. FOXM1-chDNA2 (chr9): forward, 5’-CTG CAA AAT GGA CCA ATC AGC-3’; reverse, 5’-AAG ATG GTG TGT CCG GAA TTT G-3’. FOXM1-chDNA3 (chr4): forward, 5’-TCA CAA GCC CCC TGT AGG-3’; reverse, 5’-TCC AAC TCT TGC CTG GTC TC-3’.

#### DNA Fluorescence *in situ* hybridization (FISH)

DNA-FISH probes were constructed by Exon Biotechnology Inc (Guangzhou, China). The probes covered genomic regions (hg38) used in this study was as follow: chr18: 103010-113570 (chr18_FOXM1-chDNA1_10 Kb). For fluorescence *in situ* hybridization assay, DNA FISH was performed as described^66^. Briefly, coverslips containing fixed and permeabilized cells were quenched by 3% H_2_O_2_, followed by dehydration in 70%, 85%, and 100% ethanol for 1 min each. FISH probe (chr18_ FOXM1-chDNA1_10 Kb) was added onto a glass slide, and lower the coverslips onto the slide slowly. Place the slides with samples to be hybridized in a heating block at 85°C for 5min, and then 37°C for 20 hours. The slide was then removed, and the sample was washed with Wash Buffer (1 X, preheated to 73°C) for 2 min. Added blocking buffer onto the hybridized samples and incubated at 37°C for 30 min. Added primary antibody onto coverslip and incubated at 37°C for 30 min. The sample was washed with Wash Buffer (1 X, preheated to 45°C) for 5 min. Added secondary antibody onto coverslip and incubated at 37°C for 30 min. Wash coverslip in Wash Buffer (1x) three times at 45°C for 5 min. Re-wash once with PBS. Preparation of TSA solution at TSA: 0.15%H2O2: TSA amplification buffer = 1:5:500-1:10:1000. Add TSA solution to coverslip and incubate 5-15min at RT. Wash coverslip in Wash Buffer (1 X) three times for 30 min. Mount coverslip with Anti-fade Fluorescence Mounting Medium (with DAPI, Abcam, #ab104139).

#### CRISPR/Cas9-mediated knock-in of a TetO array (96x) for visualization of FOXM1-chDNAs

To obtain a TetO-knockin cell line, we designed a pair of gRNAs using http://crispr.mit.edu/. The pair of gRNAs (5’-CAC CTG GGT GAT CAG AGC AA-3’ and 5’-TCT AGG CTT TGG CCT ACA GG-3’) bound to FOXM1-chDNA1 or gRNAs (5’-GCA GGC AGA GCG TAA GCA AA-3’ and 5’-GGA CAC ACG TTT AAT CGA GT-3’) bound to FOXM1-chDNA3 was cloned into the pX335 vector (Addgene, 42335) before the gRNA scaffold. Furthermore, we designed two donor plasmids that contain the TetO array (96x) flanked by 500-bp homology arms of the FOXM1-chDNAs in the pSP2-96-merTetO-EFS-BLaR vector (Addgene, 118713). The primers (5’-CCC TTT CGT CTT CAA GAA TTC CAT CAC CTG GGT GAT CAG TGT AGA-3’, 5’-ACC CAT TCC TAG GGC GAA TTC GCT CTC TGA TCA CCC AGG TGA T-3’; 5’-GCG CTGC TAG CTT AAG GTA CCA GCG TAG GCC AAA GCC TAG AC-3’, 5’-CCC AGA TCT ATC GAT GGT ACC CTT GTC TAG GAT CTG CCT ACA GGG-3’) were synthesized and amplified homology arms of the FOXM1-chDNA1. The primers (5’-CCC TTT CGT CTT CAA GAA TTC CCT GTA GGC AAG CCT ACA CAA GT-3’, 5’-ACC CAT TCC TAG GGC GAA TTC AGC GCT TAC GCT CTG CCT G-3’; 5’-GCG CTG CTA GCT TAA GGT ACC GAC CGA TTA AAC GTG TGT CCT TT-3’, 5’-CCC AGA TCT ATC GAT GGT ACC GTT TTT TCC TTT AAG ACT TAT GTA ATG AAT T-3’) were synthesized and amplified homology arms of the FOXM1-chDNA3. The paired gRNAs and donor plasmids were co-transfected into A549 cells. Blasticidin (10 μg/ml) selection was started 1 day after transfection. Three weeks after blasticidin selection, TetO-knockin cell clones were transplanted into 24-well plates. Cells were collected from 24-well plates and extracted genomic DNA (gDNA) for genotyping.

For FOXM1-chDNAs loci visualization, the TetO-knockin cells were transiently expressed with TetR-EGFP. Cells were fixed in 4% paraformaldehyde and mounted with Mounting Medium with DAPI (Abcam, ab104139). Images were captured by FV1200 laser scanning microscopes (Olympus, Japan). For qPCR, the following primers were used for qPCR analyses of knock-in of FOXM1-chDNAs. TetO-Telomere DNA (FOXM1-chDNA1 locus): forward, 5’-GAA GAC TAC AGC GTC GCC AG-3’; reverse, 5’-CGC GAC GAT ACA AGT CAG GT-3’. TetO-DUX4 DNA (FOXM1-chDNA3 locus): forward, 5’-TCT GAA GAC TAC AGC GTC GC-3’; reverse, 5’-ACA CAT AAC CAG AGG GCA GC-3’.

#### Extracellular vesicle isolation

Cells seeded in 150 mm culture dishes at approximately 80% confluence were incubated with serum-free DMEM for 1 day or 2 days. Cell viability was assessed using trypan blue and only > 95% viability was used for EVs isolation. The cell-conditioned medium was collected by sequential ultracentrifugation^1,18^. Briefly, the medium was first subjected to a centrifugation at 400 × g for 10 min to remove cells. Next, the supernatant was centrifuged at 2,000 × g for 20 min to remove debris and apoptotic bodies. Then, the supernatant was centrifuged at 15,000 × g for 40 min to obtain large EVs. The resulting large EVs pellet was resuspended in a large volume of PBS buffer followed by ultracentrifugation at 15,000 × g for 40 min to wash the sample. To remove any remaining any large EVs, the media supernatant from the first 15,000 × g step was passed through a 0.22-μm pore PES filter (Millipore, Bedford, MA, USA). This supernatant (pre-cleared medium) was next subjected to ultracentrifugation at 120,000 × g for 4 h in a SW 28 Ti Swinging-Bucket Rotor (Beckman Coulter, Fullerton, CA) to sediment small EVs. The crude small EVs pellet was resuspended in a large volume of PBS followed by ultracentrifugation at 120,000 × g for 4 h to wash the sample. All following centrifugation steps were performed at 4°C. Notably, for the comparison of EVs in different conditions, the results must be corrected by total cell number or total protein concentration to eliminate differences in cell seeding.

## Acknowledgements

We thank Dr. Zhuoxian Rong of Central South University for help with construction of FOXM1 knockout A549 cell line by CRISPR/Cas9 technology; thank Dr. Jianhui Jiang of Hunan University for help with confocal microscopy. This work was supported by the National Natural Science Foundation of China (grant number 81773169 to Y.T.), China Changsha Development and Reform Commission “Mass entrepreneurship and innovation program” (2018-68) and “Innovation platform construction program” (2018-216).

## Author contributions

Y.Z. and Y.T. conceived this project. Y.Z., N.D., Y.L., X.Z., S.Y., H.L. and Y.W. performed all of experiments. Y.Z., H.C., P.F., Y.P., L.Y. and T.Y. analyzed ChIP-seq and evDNA-seq. M.O. performed molecular docking simulation. Y.C, L.Y. and X.H. provided administrative and material supports. Y.T. supervised this study. Y.Z. and Y.T. composed the manuscript. All authors reviewed the manuscript and discussed the work.

## Supplementary Material

Supplementary methods, figures and tables.

