## Supplementary Materials for "Transcription factor FOXM1 specifies the loading of chromatin DNA to extracellular vesicles"

**Supplementary Figures**

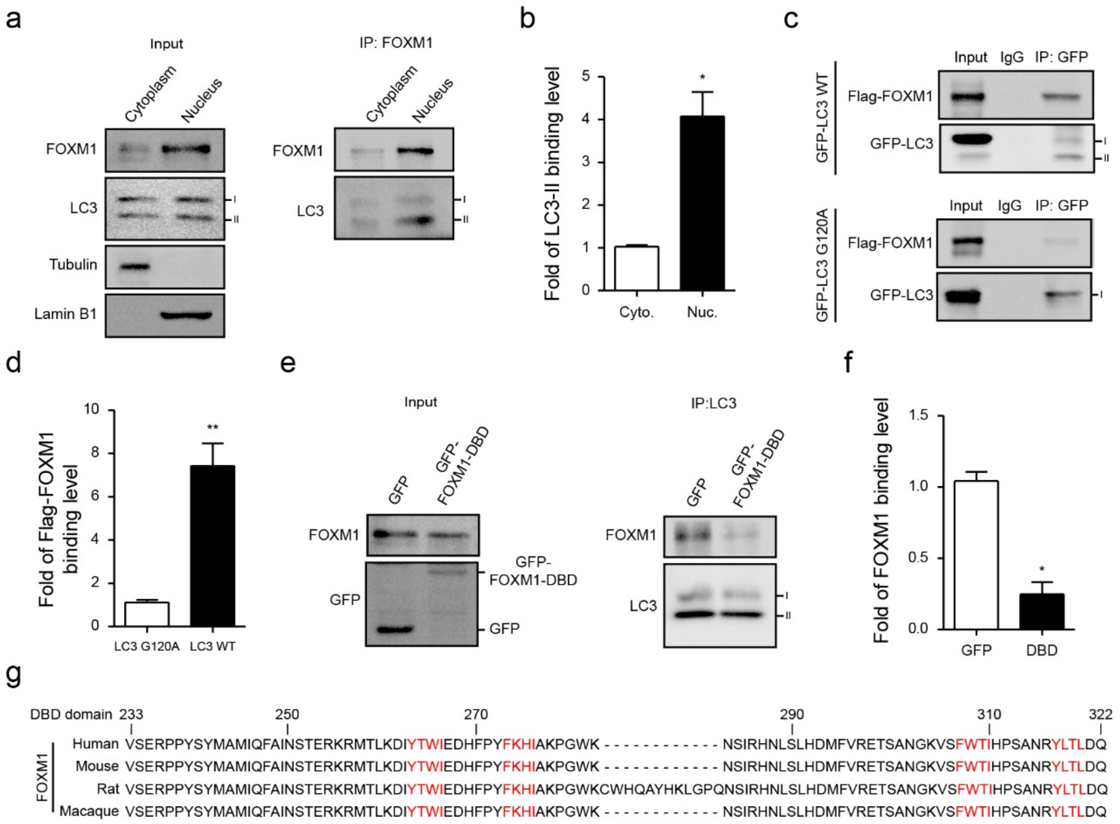


**Extended Data Figure 1. FOXM1-LC3 interaction. (a)** Nuclear and cytoplasmic components were extracted from A549 cells. All components were immunoprecipitated by anti-FOXM1 antibodies and subjected to immunoblotting. **(b)** Quantification of LC3-II binding levels in cytoplasm and nucleus (n = 3 independent experiments). **P*<0.05, unpaired two-tailed**. (c)** HEK293T cells were co-transfected with Flag-FOXM1 and GFP-LC3 WT or G120A lipidation-deficient mutant constructs and subjected to GFP immunoprecipitations. **(d)** Quantification of Flag-FOXM1 binding levels in GFP-LC3 WT and G120A mutant samples (n = 3 independent experiments). ***P*<0.01, unpaired two-tailed**. (e)** A549 cells were transfected with GFP or GFP-FOXM1-DBD constructs and subjected to LC3 immunoprecipitations. **(f)** Quantification of FOXM1 binding levels in GFP and GFP-FOXM1-DBD samples (n = 3 independent experiments). **P*<0.05, unpaired two-tailed**. (g)** Comparison of FOXM1 DBD domain amino acid sequences in different species (human, mouse, rat and macaque).

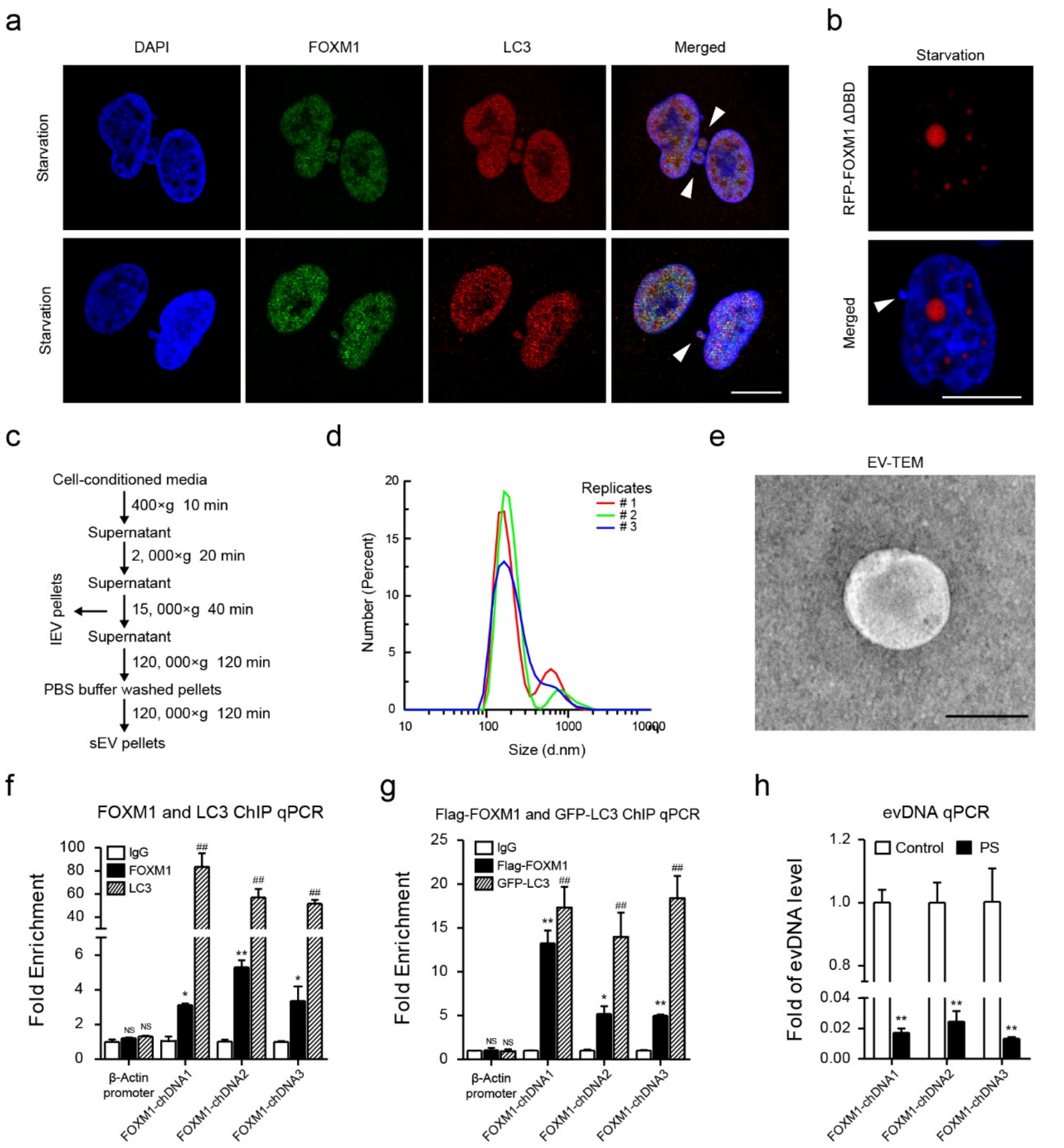


**Extended Data Figure 2. The Characterization of FOXM1-chDNAs in extracellular vesicles. (a)** The LC3-FOXM1-DNA vesiculation in cytoplasm of starved A549 cells (6 hrs). The white arrows indicated LC3-FOXM1-DNA vesicle. **(b)** HEK293T cells were transfected with RFP-FOXM1 ΔDBD (DBD domain deleted mutant) plasmid (1 μg, 24 hrs). Representative RFP-FOXM1 ΔDBD images in starved cells (6 hrs) were shown. The white arrows indicated chromatin DNA vesicle that did not contain RFP-FOXM1 ΔDBD signal. Scale bar, 10 μm. **(c)** Centrifugation protocol and workflow for EVs enrichment. **(d)** Representative nanoparticle size analysis of EVs. **(e)** Transmission electron microscopy imaging of EVs. Scale bar, 200 nm. **(f)** A549 cells were starved (6 hrs) and subjected to FOXM1 or LC3

ChIP assays. The qPCR analyses were performed for the three FOXM1-chDNAs in DNA samples purified from FOXM1 or LC3 ChIP. **(g)** HEK293T cells were transfected with Flag-FOXM1 or GFP-LC3 plasmids (10μg, 24 hrs). All cells were starved (6 hrs) and subjected to Flag-FOXM1 or GFP-LC3 ChIP assays. The qPCR analyses were performed for the three FOXM1-chDNAs in DNA samples purified from Flag-FOXM1 or GFP-LC3 ChIP. **(h)** The qPCR analyses of FOXM1-chDNAs in the Plasmid-Safe ATP-dependent DNase (PS)-digested EVs collected from starved A549 cells (48 hrs). The Bars, mean±s.e.m.; n=3; **P*<0.05, ***P*<0.01, ##*P*<0.01; NS, non-significant; unpaired two-tailed.

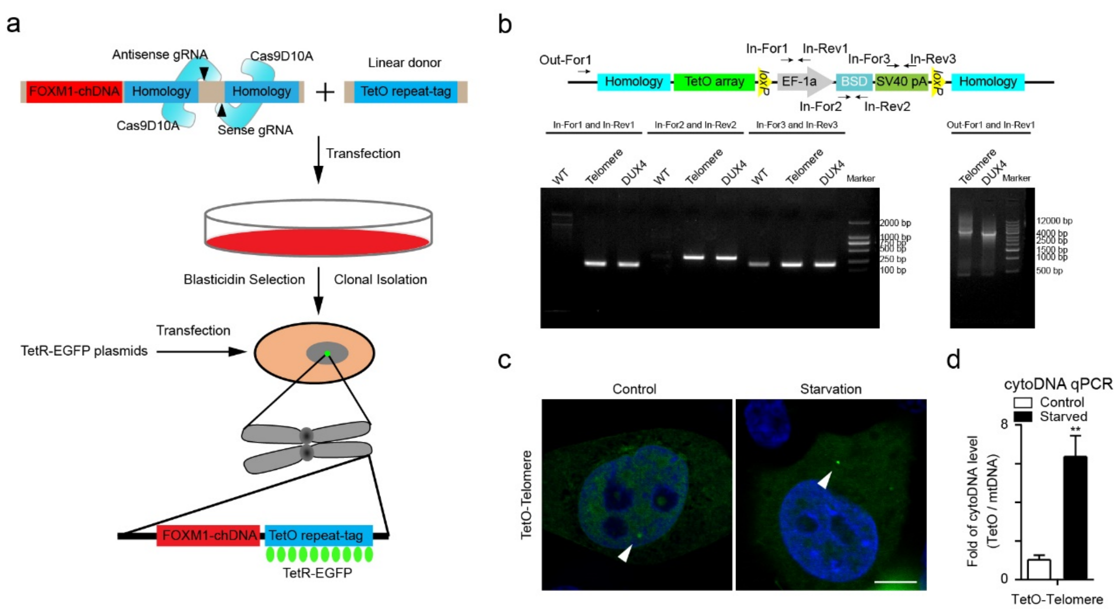


**Extended Data Figure 3. The verification of TetO-tagged FOXM1-chDNAs in cells. (a)** Overview of the workflow for the imaging of FOXM1-chDNA loci via the knock-in of a TetO array (96x) and the expression of TetR-EGFP. **(b)** The confirmation of the TetO array (96x) knock-in cells. Schematic of the primers used for genotyping was shown on top. In-For and In-Rev primers were used for the PCR analysis of A549 wild-type (WT) or the TetO array (96x) knock-in cell genomic DNA. Out-For1 and In-Rev1 primers were used for the PCR analysis of $A549^{TetO\text{-}Telomere}$ and $A549^{TetO\text{-}DUX4}$ cell genomic DNA. **(c)** Representative TetO-Telomere images in starved $A549^{TetO\text{-}Telomere}$ cells (6 hrs) that were exogenously expressed TetR-EGFP. The white arrows indicated TetO-Telomere. Scale bar, 10 µm. **(d)** The $A549^{TetO\text{-}Telomere}$ cells were treated with starvation and cytoplasmic DNA (cytoDNA) was extracted 24 hours later for qPCR. The mtDNA was internal control. The Bars, mean±s.e.m.; n=3; ***P*<0.01; NS, non-significant; unpaired two-tailed.

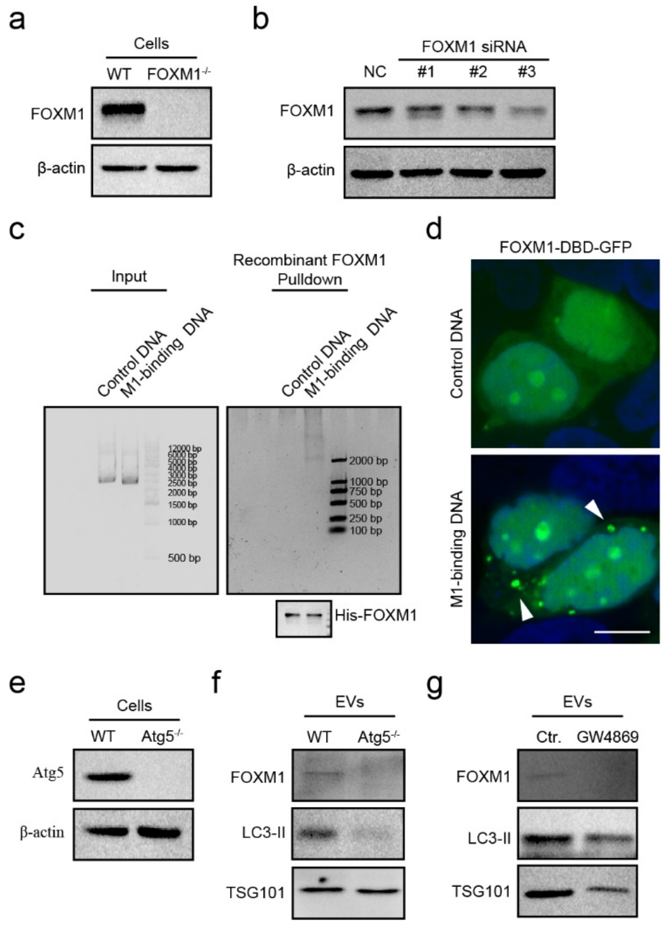


**Extended Data Figure 4. The mechanisms of FOXM1-chDNAs loading to EVs. (a)** The $A549^{FOXM1-/-}$ cell line was generated by the CRISPR/cas9 technology targeting *FOXM1* gene in A549 cells. The A549 and $A549^{FOXM1-/-}$ cell lysates were blotted by anti-FOXM1 antibodies. **(b)** The A549 cells were transfected with synthetic *FOXM1* siRNA#1, #2, #3, or with negative control (NC) siRNA (50 nM, 48 hrs). All cell lysates were blotted by anti-FOXM1 antibodies. β-actin was used as a loading control. **(c)** The linear M1-binding or Control DNA (20 μg) were incubated with recombinant protein His-FOXM1 protein (50 μg) *in vitro*. The bound protein-DNA complexes were pulled down with His-tag Resin and subjected to vertical gel electrophoresis or immunoblotting. **(d)** Representative FOXM1-DBD-GFP images in A549 cells that were co-transfected with linear M1-binding or Control DNA (1 μg) and FOXM1-DBD-GFP expression plasmid (1 μg). The white arrows indicated the complexes of FOXM1-DBD-GFP and M1-binding DNA *in vivo*. Scale bar, 10 μm. **(e)** The $A549^{Atg5-/-}$ cell line was generated by the CRISPR/cas9 technology targeting *Atg5* gene in A549 cells. The A549 and A549$^{Atg5-/-}$ cell lysates were blotted by anti-Atg5 antibodies. β-actin was used as a loading control. **(f)** The EVs were collected from starved A549 or A549$^{Atg5-/-}$ cells (48 hrs). The lysates of the EVs were blotted by anti-FOXM1 or anti-Atg5 antibodies. TSG101 was used as a loading control. **(g)** The EVs were collected from the equal numbers of starved A549 cells that were non-treated or treated with GW4869 (20 μM, 24 hrs). The lysates of the EVs were blotted by anti-FOXM1, anti-Atg5, and anti-TSG101 antibodies.

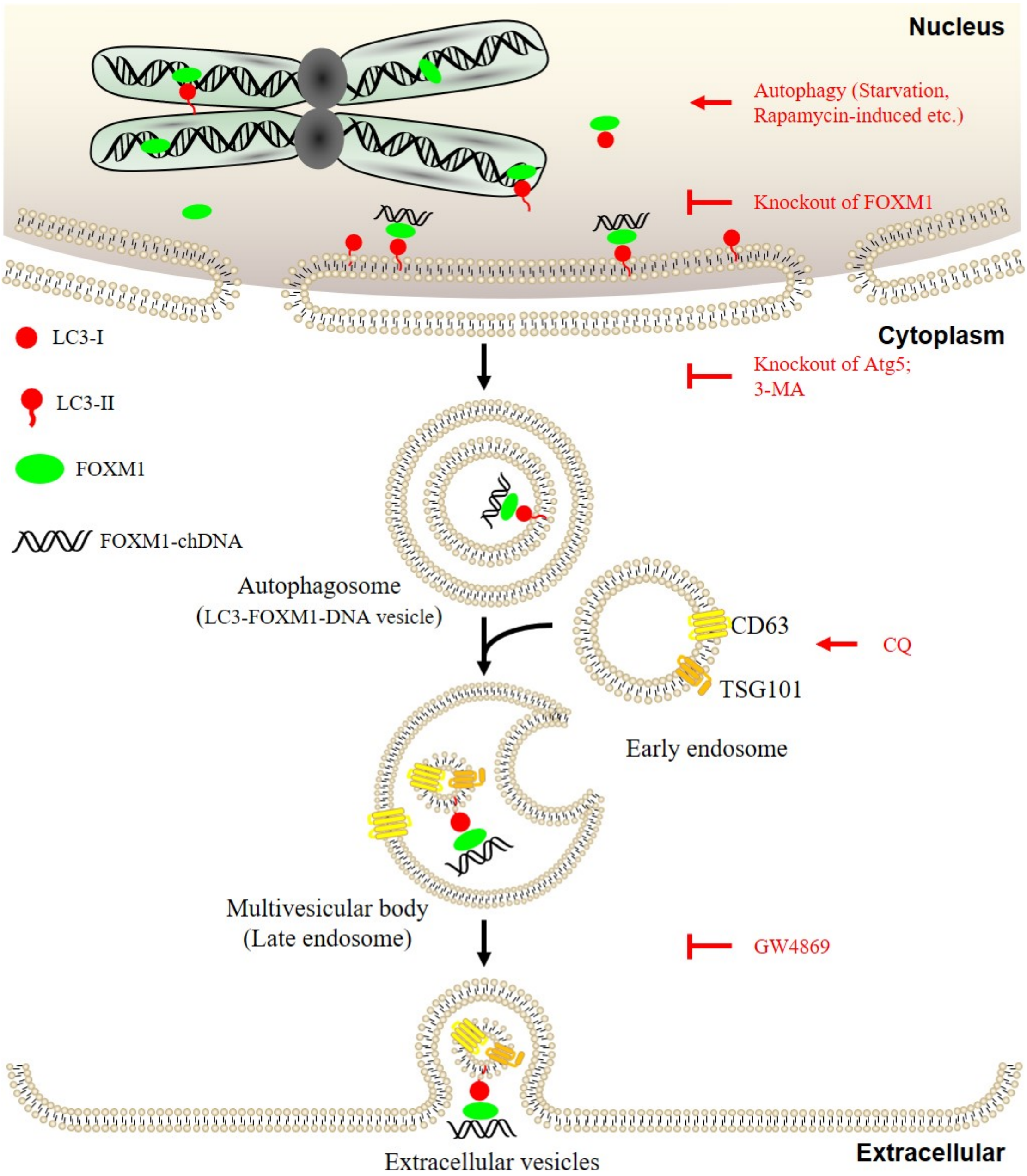


**Extended Data Figure 5. Schematic illustration of the loading of FOXM1-chDNAs to EVs.** Atg5[1], autophagy regulator essential for LC3-II lipidation; CD63[2] and TSG101[3], MVB specific markers; CQ, an autolysosome inhibitor[4]; GW4869, a specific inhibitor of neutral sphingomyelinase for inhibiting EVs secretion[5].

### Supplementary Tables

Table S1. FOXM1 and LC3 co-binding chromatin loci.

**Supplementary Materials and Methods**

**Cell culture.** The human lung epithelial cell line A549 (ATCC, CCL-185) and human embryonal kidney cell line HEK293T (ATCC, CRL-3216) were cultured in DMEM (Thermo Fisher, 11995123) supplemented with 10% fetal bovine serum (FBS, Thermo Fisher, 10270106), 100 U ml$^{-1}$ penicillin, and 100 μg ml$^{-1}$ streptomycin (Thermo Fisher, 15070063). For starvation, cells were cultured in Earle's Balanced Salt Solution (EBSS, Thermo Fisher, 24010043).

**Plasmid constructs.** The full length of human FOXM1 (NM_202002.3) was amplified from pcDNA3.1-Flag-FOXM1 and cloned into the pCMV-Tag3B, pEGFP-C2, pBiFC-VN173 and pcDNA3.1-RFP vectors by one-step cloning to yield Myc-FOXM1, GFP-FOXM1, VN173-FOXM1 and RFP-FOXM1, respectively. The FOXM1 point mutations and truncations were obtained from these constructs by inverse PCR and self-ligation. The prokaryotic expression of FOXM1 was cloned into pET-25b vector.

The full length of human LC3 (NM_022818.5) was amplified from pEGFP-LC3 and cloned into the pBiFC-VC155 and pGEX-4T2 vectors by one-step cloning to yield VC155-LC3 and GST-LC3, respectively. The LC3 point mutations and truncations were obtained from these constructs by inverse PCR and self-ligation. The Flag-FOXM1 and GFP-LC3 fragments were amplified and cloned into pLVX-IRES-Puro lentiviral expression vector. The linear 72x M1-binding DNA were synthesized by TSINGKE Inc (Changsha, China).

**Reagents and antibodies.** Chloroquine (CQ, C6628), 3-Methyladenine (3-MA, M9281) and Rapamycin (V900930) were purchased from Sigma-Aldrich. GW4869 (HY-19363) was purchased from MCE. The following antibodies were used: LC3 (Proteintech 146001AP for immunoprecipitation and WB; Cell Signaling Technology 3868 for ChIP, IF and WB), FOXM1 (Santa Cruz sc376471 for IF and WB; Cell

Signaling Technology 20459 for ChIP, IP, IF and WB), CD63 (Proteintech 25682-1-AP), DYKDDDDK Tag (Cell Signaling Technology 14793 and Beyotime AF0036), Myc-tag (Cell Signaling Technology 2276 and 2278), GFP-tag (Beyotime AG279 and Abcam ab290), GFP-Trap (Chromotek gta20), Lamin B1 (Beyotime AF1408), β-tubulin (Beyotime AT809), GST (Beyotime AF2299), β-actin (Sigma-Aldrich A5441), TSG101 (Beyotime AF8259), Rab7 (Beyotime AF2458).

**Lentiviral packaging, infection and selection.** To package lentivirus, HEK293T cells were co-transfected with the packaging vectors psPAX2 and pMD.2G, lentiviral constructs (such as pLVX-GFP-LC3, pLVX-Flag-FOXM1, lentiCRISPR-FOXM1 sgRNA and lentiCRISPR-Atg5 sgRNA). Virus was filtered through a 0.45-μm filter (Millipore, Bedford, MA, USA) and mixed with Polybrene (Millipore, Bedford, MA, USA) to a final concentration of 8 μg $ml^{-1}$. Subsequently, A549 or HEK293T cells were incubated with viral mix for 24 h and selected with 1 μg $ml^{-1}$ puromycin for about 1 week.

**GST-pulldown assay.** HEK293T cells (5 x $10^6$) were transfected with 5 μg plasmids (such as Myc-tagged FOXM1 and mutations) for 24 hrs and were lysed in the IP lysis buffer (50mM Tris-HCl pH 7.4, 150mM NaCl, 1% NP-40, 5mM EDTA, 5mM EGTA, 5% Glycerol, containing 1 × protease inhibitor cocktail) and centrifuged to obtain the supernatants. The supernatants were incubated with 30 μg bacterially purified GST-LC3 or truncated proteins and additional 30 μl GST beads (washed three times with PBS buffer) for 2 h at 4°C. The complex was washed 5 times with PBS buffer (containing 1 mM phenylmethylsulphonyl fluoride, PMSF). The proteins were eluted with 1 × loading buffer by boiling for 10 min and analyzed by 10% SDS-PAGE followed by immunoblotting with anti-Myc antibody.

**Immunofluorescence staining**. Cells were treated as indicated and fixed in 4% paraformaldehyde in PBS for 40 min at 4°C, washed twice with TXBST (0.1% Triton X-100 in TBS). Cells were then blocked in 5% BSA in TBS for 1h at room temperature and incubated with primary antibodies in 5% BSA in TBS supplemented with 0.1% Tween 20 (TBST) overnight at 4℃. Cells were washed 3 times with TBST, each for 5 min, followed by incubation with Alexa Fluor-conjugated secondary antibodies (Abcam) in TBST (containing 5% BSA) for 1h at room temperature. Cells were then washed 3 times in TBST and mounted with Mounting Medium with DAPI (Abcam, ab104139). Images were captured by FV1200 laser scanning microscopes (Olympus, Japan).

**Immunoprecipitation.** Cells ($4 \times 10^7$) were lysed in the IP lysis buffer and centrifuged to obtain the supernatants. The supernatants were rotated with anti-IgG antibody (Sigma-Aldrich, 12-370) and additional Protein A/G magnetic beads (Thermo Scientific, 88802) for 4h at 4℃. The supernatants were collected and incubated with antibody-conjugated Protein A/G magnetic beads, and rotated at 4℃ overnight. The immunoprecipitation was washed five times with PBS buffer (containing 1mM PMSF), and boiled with 1 × loading buffer for 10 min. All samples were analyzed by western blotting.

**Molecular docking.** LC3 protein was obtained from PDB ID 2N9X and FOXM1-DBD was obtained from PDB ID 3G73. We confirmed that the LIR motif (317-320aa) of FOXM1-DBD and F52 of LC3 mediated the interaction of FOXM1 and LC3. Therefore, ClusPro[6] (jobID=701602) was first used for rigid docking under restricted conditions. After that, we separated the top1 structure to LC3 and FOXM1-DBD and use Rosetta Relax Application[7] to build the ensembles of LC3 and FOXM1-DBD. Then RosettaDock[8] was used to conduct flexible docking between the two ensembles for 10,000 times.

**Nuclear and cytoplasmic components extract.** Cells ($4 \times 10^7$) were washed with PBS buffer and added 5 pellet volumes of CE buffer (10 mM HEPES pH 7.6, 60 mM KCl, 1 mM EDTA, 0.075% NP-40, 1 mM DTT and 1 mM PMSF) to cell pellet (approximately 100 μL). Incubated on ice for 3 min. Spined the preparation using a microcentrifuge at 1000-1500 rpm for 4 min. Removed the cytoplasmic extract from the pellet to a clean tube. Washed the nuclei with 100 μL of CE buffer without detergent. Be careful to resuspend the fragile nuclei gently. Spined the nuclei as above at 1000-1500 rpms for 4 min. Added 1 pellet volume NE buffer (20 mM Tris-HCl pH 7.5, 420 mM NaCl, 1.5 mM $MgCl_2$, 0.2 mM EDTA, 1 mM PMSF and 25% Glycerol) to nuclear pellet (approximately 50μL). Adjust the salt concentration to 400 mM using 5 M NaCl (add ~35 μL). Add an additional pellet volume of NE buffer. Vortex to resuspend the pellet. Incubate the extract on ice for 10 minutes. Vortex the mixture periodically to resuspend the pellet. Spin the CE and NE at maximum speed for 10 minutes to pellet any nuclei. Transfer the contents of the CE tube and NE tube separately to clean tubes. Add glycerol to the CE tube to 20%. Store at -70°C.

**Recombinant protein expression and purification.** All prokaryotic expression plasmids (pGEX-4T2 or pET-15b) were transformed into *Escherichia coli* BL21(DE3), inoculated in LB medium (containing Amp) and growth at 37°C until an optical density at 600 nm ($OD_{600}$) of 0.6 was reached, then isopropyl-β-D-thiogalactopyranoside (to a final concentration of 0.8 mM) was added. After growing for 12 h at 16°C, cells were pelleted and lysed in PBS buffer (containing 1 × protease inhibitor cocktail and 1mM PMSF). The lysate was sonicated and centrifuged to obtain recombinant protein supernatants. The GST or GST-fused recombinant proteins were purified by BeyoGold™ GST-tag Purification Resin[9,10] (Beyotime, P2250, Beijing, China) following the manufacturer's instruction. The His-fused recombinant proteins were purified by BeyoGold™ His-tag Purification Resin (Beyotime, P2233, Beijing, China) following the manufacturer's instruction.

**Chromatin immunoprecipitation (ChIP) assay.** For FOXM1 or Flag-FOXM1 ChIP, cells ($4 \times 10^7$) were crosslinked by formaldehyde and stopped by glycine. Cells were scraped and lysed with Nuclear

Lysis Buffer (1% SDS, 10 mM EDTA, 50 mM Tris, pH 8.0, protease inhibitor) for 10 min on ice. The resulting extract was sonicated and taken 20% of samples as Input control. Other samples were performed immunoprecipitation. For LC3 or GFP-LC3, ChIP was performed as previously described[11]. All samples were reversed crosslinks and purified DNA to qPCR or sequence.

**Isolation of MVBs by sucrose density gradient centrifugation.** For the separation of MVBs, cells (4 × $10^7$), cultured in DMEM with 10% FBS, were washed with ice-cold PBS and collected with a cell scraper. Centrifuge the cells at 300g for 5 min at 4°C. Cell pellet was loosened using homogenization buffer [HB; 250 mM sucrose, 3 mM imidazole (pH 7.4), 1 mM EDTA, 0.03 mM cycloheximide, 10 mM iodoacetamide, 2 mM phenylmethylsulfonyl fluoride (PMSF), and 1× cocktail inhibitor]. Centrifuge at 1300g for 10 min at 4°C. The supernatants were discarded, and the pellet was gently resuspended with a wide-cut tip in three times the pellet volume of HB. The suspension was passed through a 1-ml syringe two to 10 times. Dilute the homogenate in HB (1 part homogenate to 0.7 parts HB). Centrifuge at 2000g for 10 min at 4°C. Collect the supernatants and centrifuge again. Carefully collect the supernatants (termed the post-nuclear supernatants [PNS]) from the second centrifugation.

Sucrose density solutions were prepared before use to generate discontinuous step (10%-62%) gradients. The PNS was carefully added to the top of sucrose density gradients (10%-62%) in a centrifugaton tube. The samples were subjected to ultracentrifugation at 140, 000 g for 15 h at 4°C using a SW28 TI Swinging Bucket rotor (*k* factor of 246, Beckman Coulter). Five individual fractions of 1 mL were collected from the top of the gradient. For immunoblotting, each individual 1 mL fraction was transferred to new ultracentrifugation tubes, diluted 25-fold in PBS and subjected to ultracentrifugation at 140, 000 g for 4 h at 4°C using a SW28 TI Swinging Bucket rotor. The resulting pellets were lysed in cell lysis buffer for 10 min on ice. For DNA extraction, the resulting pellets were extracted by QIAamp

DNA Mini Kit (Qiagen, Valencia, CA, USA) following the manufacturer's instruction.

**Rapid method for the immune-purification of MVBs (IP-MVBs).** Cells were quickly washed twice with PBS and then scraped in one ml of KPBS (130 mM KCl, 10 mM KH2PO4, pH 7.25 was adjusted with KOH) and centrifuged at 1, 000 g for 2 min at 4°C. Pelleted cells were resuspended in 950 μL, and 25 μL (equivalent to 2.5% of the total cells) was reserved for further processing of the whole-cell fraction. The remaining cells were gently homogenized with 20 strokes of a 2 ml homogenizer. The homogenate was centrifuged at 1, 000 g for 2 min at 4°C. The magnetic beads were incubated with anti-CD63 antibody for 1 hour. Subsequently, the supernatants were incubated with the magnetic beads for an additional hour at 4°C. Immunoprecipitates were gently washed three times with KPBS on a magnetic rack. For immunoblotting, the samples were lysed with IP buffer and analyzed by SDS-PAGE. For DNA extraction, the samples were extracted by QIAamp DNA Mini Kit (Qiagen, Valencia, CA, USA) following the manufacturer's instruction.

**Protein and DNA extraction from extracellular vesicles.** The EV pellets were lysed by urea lysis buffer (100 mM $NaH_2PO_4$, 10 mM Tris-HCl pH 8.0, 8 M Urea, 10 mM Imidazole, containing 1 × protease inhibitor cocktail) and analyzed by immunoblotting[12]. The extracellular vesicle DNAs (evDNAs) were extracted by QIAamp DNA Mini Kit (Qiagen, Valencia, CA, USA) following the manufacturer's instruction.

**Direct immunoaffinity capture (DIC) of EVs.** DIC assays of EVs were performed as described[13]. Briefly, the cell-conditioned medium was collected as described above. After ultracentrifugation at 15,000 × g for 40 min, the supernatants were incubated with magnetic beads directly conjugated to FOXM1 antibody or GFP-Trap beads for 16 h. After incubation, all beads were washed for immunoblotting or evDNAs extraction.

**Biotinylated DNA pull-down assay.** For the biotinylation of FOXM1-chDNA3 (chr4: 190179246-190180026), the 979 bp biotinylated DUX4 DNA was prepared by PCR amplification using a pair of 5'biotinylated primers: 5'-biotin-ACT CCA CTC CGC GGA GAA-3' and 5'-biotin-CTT GTC AAG GTT TGG CTT ATA GG-3'. The PCR products were purified by Universal DNA Purification Kit (TIANGEN, DP214, Beijing, China) following the manufacturer's instruction.

For DNA-pulldown, A549 cells were lysed in IP lysis buffer and centrifuged to obtain the supernatants. The supernatants were incubated with 500 ng biotinylated DUX4 DNA for 1 h and additional 50 μl streptavidin beads for another 1 h at room temperature. The complex was washed 5 times with IP lysis buffer. The proteins were eluted with 1 × loading buffer by boiling for 10 min and analyzed by 10% SDS-PAGE followed by immunoblotting with primary antibodies[14].

**Electrophoretic mobility shift assays (EMSA).** EMSA was performed as previously described[15]. Briefly, His-FOXM1 protein (10 μg) were incubated with the FAM-labeled DNA probe (50 nM) containing putative FOXM1 sites in binding buffer (20 mM Tris-Cl, 50 mM KCl, 10% glycerol, 0.5 mM EDTA, 0.2 mM DTT, pH 7.6) for 30 min on ice. The purified GST-LC3 protein was used to detect supershift.

**CRISPR/Cas9-mediated knock-out of *FOXM1* and *Atg5*.** To obtain A549$^{FOXM1-/-}$ or A549$^{Atg5-/-}$ cell line, we designed their sgRNA using http://crispr.mit.edu/. The sgRNA (5'-GAT GGC CAC TAC TTG CGT GT-3') targeted to *FOXM1* or sgRNA (5'-GAT GGA CAG TTG CAC ACA CT-3') targeted to *Atg5* was cloned into the lentiCRISPR v2 vector (Addgene plasmid #52961). FOXM1 sgRNA or Atg5 sgRNA lentivirus were packaged and transduced to A549 cells. The transduced cells were selected with 1 μg ml$^{-}$

$^{1}$ puromycin for 2 weeks to obtain the FOXM1$^{-/-}$ or Atg5$^{-/-}$ cells.

**siRNA-mediated silencing.** All siRNA were designed and synthesized by GenePharma (Shanghai, China). *FOXM1* siRNA was synthesized as follows: #1 5'-UUU CAC UUG GGG CAU UUU GAA-3'; #2 5'-UGG UUA AUA AUC UUG AUC CCA-3' and #3 5'-GGA CCA CUU UCC CUA CUU UTT-3'. Negative control: 5'-UUC UCC GAA CGU GUC ACG UTT-3'. For *FOXM1* silencing, cells were transiently transfected with siRNA (50 nM) using Lipofectamine 2000 (Life Technologies, Carlsbad, CA) according to the manufacturer's protocols.

**Statistical analysis.** All the statistical analyses were performed using GraphPad Prism 5 software. Student's *t*-test was used for comparison between two groups. One-way ANOVA with Dunnet's post-test analysis was performed to evaluate differences between groups of three or more. Significance was considered when the *P*-value was less than 0.05.
